# PIVOT: an open-source tool for multi-omic spatial data registration

**DOI:** 10.1101/2025.06.08.658506

**Authors:** André Forjaz, Valentina Matos Romero, Ian Reucroft, Margaret Eminizer, Donald Kramer, Daniela Higuera, Hengameh Mojdeganlou, Paola A. Guerrero, Jimin Min, Meredith Wetzel, Dmitrijs Lvovs, Alens Valentin, Sarah M. Shin, Xuan Yuan, Rosalie C. Sears, Koei Chin, Anirban Maitra, Elana J. Fertig, Won Jin Ho, Luciane T. Kagohara, Laura D. Wood, Denis Wirtz, Dimitrios N. Sidiropoulos, Ashley L. Kiemen

## Abstract

Advances in spatial profiling have resulted in the generation of multi-omic atlases that span biological scales. In general, multiple workflows are required for image registration, coordinate registration, and spot deconvolution to integrate modalities. To improve the throughput of registration of multi-omic cohorts, we introduce PIVOT, a user-friendly and open-source interface for streamlined nonlinear registration. We demonstrate PIVOT’s strengths through registration of three multi-omic datasets, and show comparison of its performance to existing workflows.

## Main text

Technological advances have improved our ability to collect diverse spatial information from tissue at the proteomic, transcriptomic, metabolomic, and genomic level.^1–5^ These advances enable imaging tissue across ontological scales via serial sectioning.^6–9^ In tandem, advances in computing and analytic workflows have improved our ability to distill meaningful conclusions from these assays.^10,11^ Yet, challenges in integrating diverse multi-omic datasets remain.

While teams have developed isolated methods to precisely co-register specific combinations of image modalities,^12–17^ there lacks a unified method for rapid and accurate co-registration across histological images with the assistance of an intuitive graphical user interface (GUI). Many automated image registration workflows incorporate nonlinear registration to correct for local tissue warping but may be applicable to only a select number of image types (hematoxylin and eosin [H&E], immunohistochemistry [IHC], etc.) and often fail in registration of small regions of interest (ROIs) to whole slide images (WSIs). On the other hand, fiducial-point-based registration workflows are applicable to any image type and excel at ROI to WSI registration, as they rely on human detection of structures. However, due to their reliance on fiducial point selection these methods are subject to inter-user variability. In general, to integrate complex multi-omic datasets, researchers combine multiple workflows, leading to inconsistent metadata formatting and requiring complex downstream synthesis of information.

Here, we introduce PIVOT, an open-source and user-friendly interface for streamlined, semi-automated registration of multi-omic images (**Fig 1**). PIVOT streamlines the registration process by combining image and coordinate registration into a single application, clustering datasets into projects that may be reloaded and modified at any time. PIVOT combines the advantages of fiducial-point based and automated nonlinear techniques, using user-guided affine registration followed by nonlinear finetuning, resulting in high-resolution registration across image types. We compared the performance of PIVOT to ImageJ landmark correspondence (affine registration using user selected fiducial points) and CODA (automated nonlinear registration designed for brightfield images) and showed that PIVOT outperforms both (**Fig S1A**).

**Fig 1.**
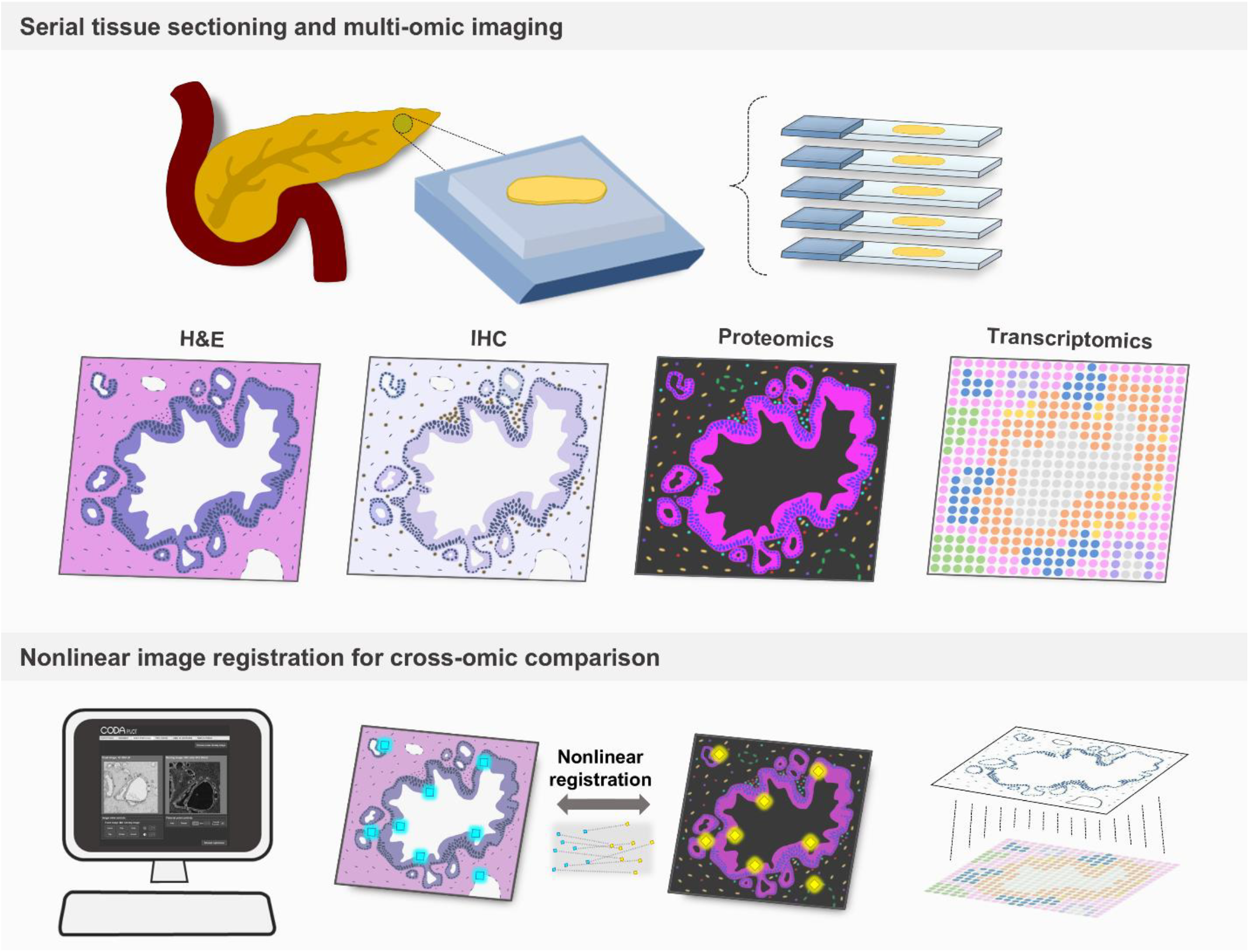
PIVOT: an open-source tool for multi-omic spatial data registration.

One weakness of fiducial point-based registration algorithms is their reliance on the quality of the point pairs selected by users, introducing user-to-user variation. To evaluate the effect of user-accuracy in fiducial placement on registration accuracy, we simulated good and poor registration conditions. Two independent users registered ten pairs of images multiple times, allowing 30 seconds, 1 minute, and 2minutes for fiducial placement (**Fig S1B**). With 30 seconds, users had little time for accurate fiducial placement, resulting in poor affine alignment. Even so, using the nonlinear finetuning, all image pairs reached the equivalent accuracies, demonstrating that user-error has minimal effect on registration quality.

To test the robustness of PIVOT across multi-omic image types, we analyzed three datasets spanning histological, molecular, proteomic, metabolomic, and transcriptomic data. We used PIVOT to register each dataset, using the high-resolution H&E image as the reference coordinate system (**Fig 2A, Fig S1C-D**). We obtained an average 2.5, 7.95, and 8.23 root-mean-squared-error (RMSE) following affine registration, and an improved 1.72, 7.47, and 5.22 RMSE following nonlinear adjustment, demonstrating the added value of the automated finetuning.

**Fig 2.**
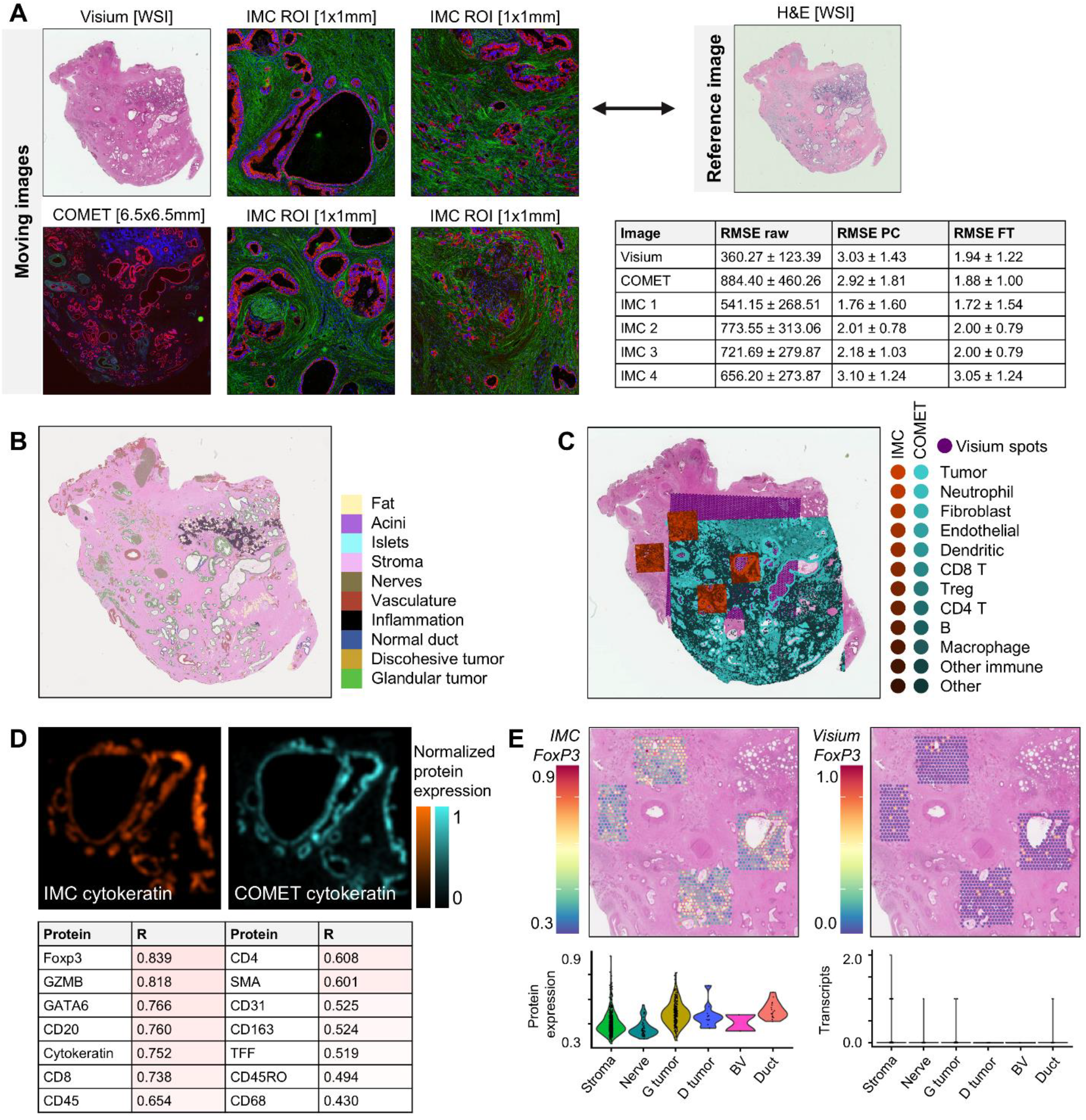
Cross-platform integration enabled by PIVOT. (**a**) Guided nonlinear registration of Visium spatial transcriptomics, COMET Lunaphore, and imaging mass cytometry (IMC) to high-resolution hematoxylin and eosin (H&E) stained human pancreas tissue. Table containing raw, affine point-cloud (PC), and nonlinear fine-tuned (FT) root-mean-squared error (RMSE). (**c**) Cell-type prediction using CODA segmentation applied to high-resolution histology. (**d**) Overlay of Visium spots (purple) with protein expression from COMET Lunaphore (cyan) and IMC (red). (**e**) Two-dimensional cross-correlation of protein expression between co-registered IMC and COMET Lunaphore shows strong agreement. (**f**) IMC-derived per spot pseudo-bulk protein expression of *FOXP3* compared to Visium-derived per spot transcript count of *FOXP3* demonstrates that poor detection of low abundance transcripts using spot-based spatial transcriptomics may be improved with proteomics. Violin plots show detection of protein and transcripts per cell-type predicted from H&E.

We applied semantic segmentation cell-type prediction to the H&E image of each dataset for detection of tumor cells, inflammation, stroma, and other histologically recognizable features (**Fig 2B, Fig S1C-D**). By registering the per-cell protein expression from COMET Lunaphore and imaging mass cytometry (IMC), and registering the spots from Visium into fixed image space, we compared protein, RNA, and histological features (**Fig 2C**). We correlated the common protein markers between COMET and IMC to show strong agreement, demonstrating the power of this tool to validate protein expression across assays (**Fig 2D**). In two additional datasets, we similarly compared the per-cell protein expression from cycIF and COMET Lunaphore to H&E-based cell type prediction, finding strong agreement (correlation coefficients ranging from 0.43 – 0.84) between immune and epithelial markers (**Fig S1B-C**).

Finally, we adapted a previously reported technique to deconvolve Visium spots using H&E-based cell type prediction to enable pseudo-bulk of protein expression from IMC data into Visium spots.^18^ We compared per-spot protein expression of key markers such as *FOXP3* to per-spot transcript reads, clustering spots by their majority cell type as determined in H&E (**Fig 2E, Fig S2**).

Expression of lymphoid cell type markers (*CD8, CD4, FOXP3, MS4A1*) showed discordance between RNA and Protein modalities consistent with gene drop out in Visium spatial transcriptomics. *CD8* protein expression was detected in ductal regions only in IMC, whereas RNA expression was primarily detected in spots with low cellularity, such as fat and stroma, suggesting poorer detection of lowly expressed transcripts in spots with high UMI counts. Similarly, regulatory T cell marker *FOXP3* was lowly expressed at the RNA level across annotations, including regions with *FOXP3* staining identified by IMC.

In conclusion, PIVOT is a robust, open-source tool for spatial alignment of multi-omic datasets, enhancing the power of spatial assays by improving researchers’ ability to compare across ontological scales.

## Data availability statement

The data analyzed here is available from the corresponding author upon request.

## Code availability statement

The PIVOT software is available for download at the following address: https://github.com/Kiemen-Lab/CODA_pivot

## Acknowledgements

The authors acknowledge the following sources of funding: BreakThrough Cancer Data Science, BreakThrough Cancer Demystifying Pancreatic Cancer Therapies, the Johns Hopkins University Data Science and Artificial Intelligence Institute (DSAI), NIHU54CA268083; Lustgarten Foundation-AACR Career development award for pancreatic cancer research in honor of Ruth Bader Ginsburg; Susan Wojcicki and Denis Troper; The Carl and Carol Nale Fund for Pancreatic Cancer Research; the Rolfe Pancreatic Cancer Foundation; Fight Cancer Stay Positive; The Sol Goldman Pancreatic Cancer Research Center. KC was supported by U2CCA233280 and Prospect Creek Foundation. JM was supported by a Seed Grant from the Hirshberg Foundation for Pancreatic Cancer Research. AM was supported by U54CA274371 and U24CA 274274. We thank Francesca Paradiso for her assistance in the generation of COMET Lunaphore data. DNK was supported by the Maryland Cigarette Restitution Fund and the Lustgarten Foundation.

## Author contributions

ALK conceived the project. HM, PG, JM, MW, DL, AV, SMS, XY, RS, KC, AM, EJF, WJH, LK, LDW, and DW generated the sample datasets. ALK led the design of the image registration application, with technical help from AF, VMR, IR, ME, DK, DH, and DS. AL wrote the first draft of the manuscript, which all authors edited and approved.

## Figures and Captions

**Fig S1.**
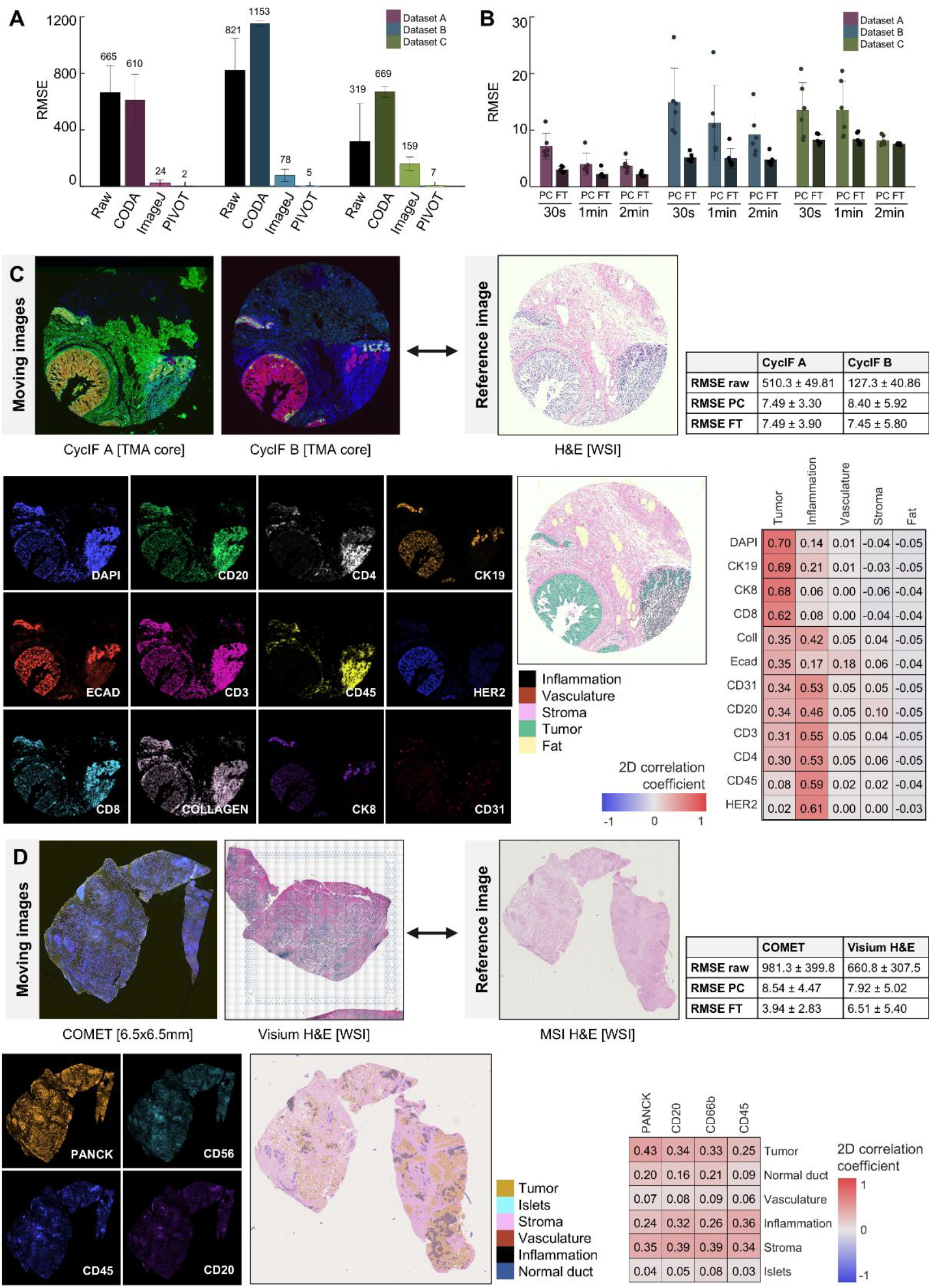
Comparison to additional datasets. (**a**) Comparison of root mean squared error (RMSE) between fiducial point pairs of unregistered images and images registered using the CODA algorithm, ImageJ landmark correspondence, and PIVOT. (**b**) Comparison of RMSE in point-cloud affine registration (PC) and finetuned nonlinear registration (FT) on three datasets allowing 30 seconds, 1 minute, or 2 minutes for fiducial placement. (**c**) Registration of cyclic immunofluorescent (cycIF) to hematoxylin and eosin (H&E), with RMSE provided. Comparison of protein expression from cycIF to cell-type predictions derived from H&E. (**d**) Registration of COMET Lunaphore and Visium to H&E, with RMSE provided. Comparison of protein expression from COMET Lunaphore to cell-type predictions derived from H&E.

**Fig S2.**
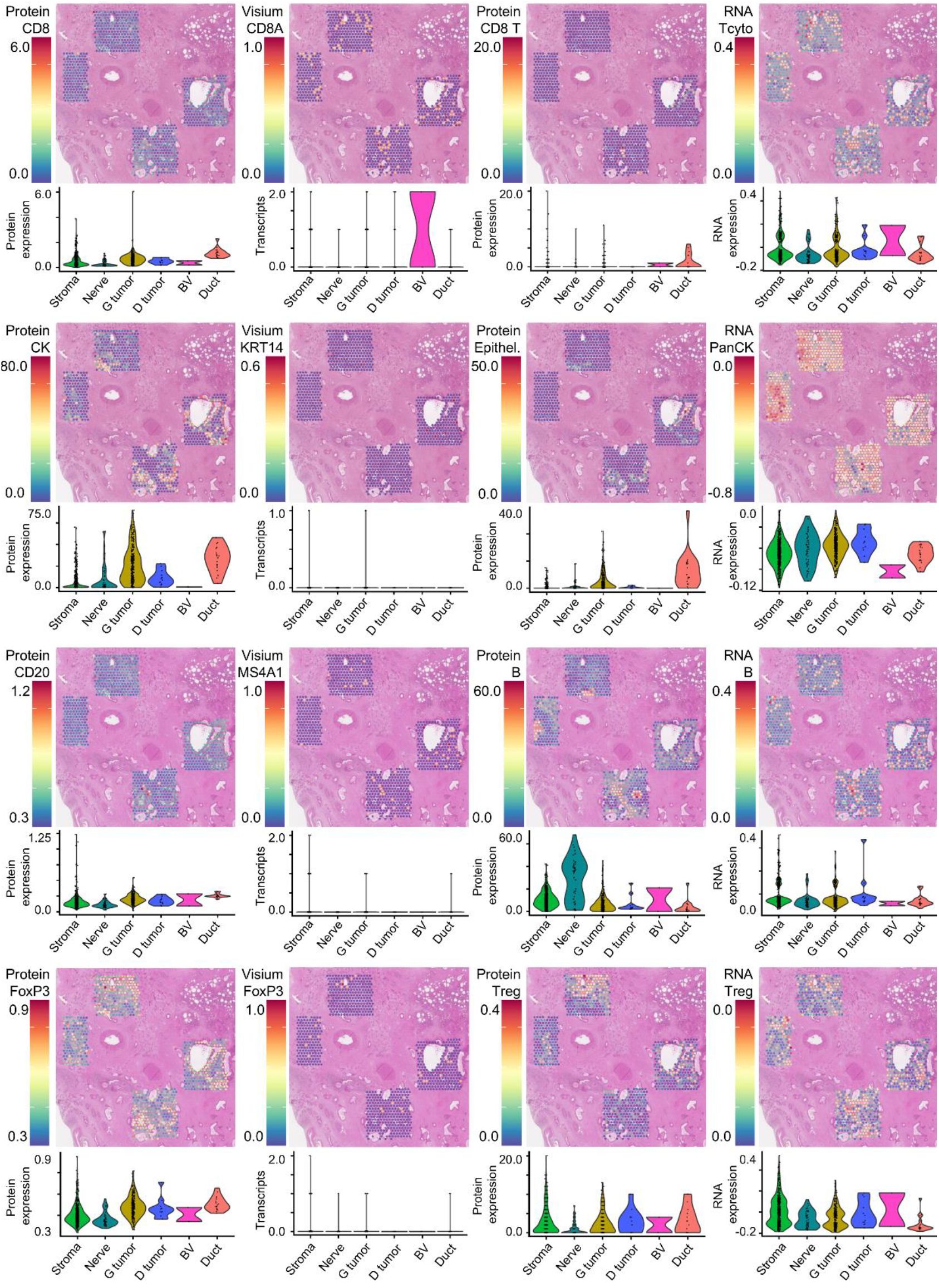
Integration of spatial transcriptomics, imaging mass cytometry, and H&E-based cell type predictions. IMC-derived per spot pseudo-bulk protein expression of several markers compared to Visium-derived per spot transcript counts and RNA expression demonstrates that poor detection of low abundance transcripts such as *FOXP3, CD8*, and *CD20* using spot-based spatial transcriptomics may be improved using proteomics. Violin plots show detection of protein and transcripts per cell-type predicted from H&E. Cytokeratin protein expression and pan-cytokeratin RNA-expression show high abundance in cells predicted as epithelial (glandular tumor, discohesive tumor, and normal duct) in H&E.

## Materials and Methods

### Generation of dataset A

A sample of human pancreatic tissue containing invasive pancreatic ductal adenocarcinoma was prospectively collected from an individual treated with neoadjuvant chemotherapy and pancreatic resection at the Johns Hopkins Hospital. Acquisition of this sample was approved by the Johns Hopkins University Institutional Review Board. The resected tissue was formalin-fixed, paraffin-embedded, and serially sectioned at a thickness of 4µm. Various spatial profiling technologies were applied to the serial sections, including: (1) hematoxylin and eosin (H&E) staining, imaged at 20x magnification using a Hamamatsu S360; (2) 10x Genomics Visium spatial transcriptomics, with one 6.5 × 6.5 mm^2^ region of interest; (3) imaging mass cytometry, with four 1 × 1 mm^2^ regions of interest; and (4) COMET Lunaphore, with one 8 × 8 mm^2^ region of interest.

### Generation of dataset B

A human breast cancer tissue microarray (TMA) was created with a series of breast cancer subtypes (Hormone Receptor Positive, HER2 Positive, Triple Negative, and Lobular), subtype associated cell lines, and normal tissues (Breast, Tonsil, and Jejunum) at Pantomics (Fairfield, CA). All samples were collected at the time of surgical resection and underwent coring, formalin-fixation, and paraffin embedding. The sample was serially sectioned. One section was stained with H&E and two sections were stained with cyclic immunofluorescence (cycIF).

### Generation of dataset C

A sample of human pancreatic tissue containing invasive pancreatic ductal adenocarcinoma was retrospectively collected from the MD Anderson Cancer Center. Acquisition of this sample was approved by the MD Anderson Institutional Review Board. The resected tissue was formalin-fixed, paraffin-embedded, and serially sectioned at a thickness of 4µm. Various spatial profiling technologies were applied to the serial sections, including: (1) hematoxylin and eosin staining, imaged at 20x magnification using a Hamamatsu S360; (2) 10x Genomics Visium spatial transcriptomics, with one 6.5 × 6.5 mm^2^ region of interest; (3) COMET Lunaphore, with one 8 × 8 mm^2^ region of interest.

### Spatial Transcriptomics

Spatial transcriptomic data was processed using the Seurat R package (v5).^5^ Visium spatial RNA-seq data (10x Genomics) was loaded with Load10X_Spatial. For each spot, the top CODA-predicted cell type was assigned based on the highest proportion of pixels present within each spot. Gene module scores were calculated using AddModuleScore for curated epithelial and immune gene signatures (*panCK: KRT* genes detected in dataset; *Tcyto: CD3E, CD3D, CD8A, GZMB; B cell: CD19, MS4A1, CD79A, CD22; NK: NCAM1, KLRD1, NCR1, PRF1, KLRK1; Treg: CD3E*,

*CD3D, CD4, FOXP3, CTLA4, PDCD1*). Raw counts of selected transcripts (e.g., *CD3E, CD8A, MS4A1, KRT14, FOXP3*) were extracted from the SCTransform assay, log-transformed, and scaled. Correlation between IMC markers and spatial transcriptomic expression was evaluated using linear regression and visualization via the ggpubr package.^19^ Spatial and violin plots were generated using Seurat functions, stratified by CODA-assigned top cell type. All outputs were compiled into a single PDF for visualization. R version 4.5.1 was used.

### Imaging Mass Cytometry

Imaging mass cytometry images were ablated using a Hyperion Imaging System (Standard BioTools). Resulting MCD files were converted into ome.tiff format using MCD Viewer software (Standard BioTools; version 1.0.560.6). These files were analyzed using the HighPlex FL (version 4.2.14) algorithm in HALO (Indica Labs; version 3.6.4134.396). The DNA1 channel was used as a nuclear marker to identify all cells. Expression of all markers within the adjacent region to the nuclear signal was averaged and recorded per cell. Algorithm parameters were manually adjusted for each marker to optimally detect signal with minimal nonspecific signal detection. The HighPlex FL algorithm allows for cell data to be stored as objects with X and Y coordinates retained to enable spatial analysis, and enabling cell-type prediction using a clustering algorithm.

### COMET Lunaphore

The COMET platform (Lunaphore Technologies) was used to capture serial immunofluorescence. FFPE tissue was baked, deparaffinized, and rehydrated, followed by autofluorescence quenching and antigen retrieval. The slide was loaded onto the COMET Stainer and processed according to manufacturer’s recommendations, undergoing iterative staining with primary antibodies, secondary antibodies, and subsequent imaging, followed by elution of the primary and secondary antibodies. After staining, imaging outputs from each cycle were automatically stitched and aligned. Images were viewed and cellular clusters were determined using the Lunaphore Viewer.

### Cyclic Immunofluorescence (cycIF)

CycIF staining of tumor tissue was completed on TNP-TMA-7 and TNP-TMA-8 using our protocol: dx.doi.org/10.17504/protocols.io.23vggn6. Antibodies used for staining are available in the previously published source data.^20^ The whole tissue core was imaged using fluorescence microscopy as described.^21^ H&E staining was performed on TNP-TMA-9. The whole tissue core was imaged using fluorescence microscopy as described.^21^

All images at full resolution and derived mask images in ome.tiff format, and all cell feature tables in three consecutive TMA tissues were generated under the funding initiatives from Human Tumor Atlas Network Phase 1 (HTAN; https://humantumoratlas.org/). All data will be fully released in public through the NCI-recognized repository: Cancer Data Service (CDS) and Seven Bridges cloud platform (SB-CGC) at Cancer Genomics Cloud (https://www.cancergenomicscloud.org/) with associated identifies: HTAN TNP-TMA, OHSU_TMA1_XXX-YY, where XXX and YY represents TMA section ID and core ID, respectively.

### Mass Spectrometry Imaging (MSI)

Tissue samples were sprayed with a solution of 10 mg/mL of 9-Aminoacridine in 70% methanol using the HTX M5 Sprayer (HTX imaging). MALDI-MS imaging was conducted on a MALDI Synapt G2-Si (Waters, USA) at 60 µm spatial resolution. After MALDI-MS imaging, the same and an adjacent tissue section were stained with H&E for pathological annotation.

### Generation of standard-format downsampled images for PIVOT registration

The PIVOT interface can read several image types including TIF, JPG, and PNG, but runs most efficiently on modestly sized images (<500 MB per file). In contrast, multi-omic images are very high-resolution with file sizes ranging from one to hundreds of GB, and come in many formats including NDPI, TIFF, and OME. To register very large image files, we suggest calculating registration transforms on downsampled copies for faster runtimes. We created a GROOVY script for streamlined down sampling of diverse image types using QuPath.^22^ QuPath is a powerful program that can efficiently read many image formats, and enables java-based programming of custom functions. Our GROOVY script helps users downsample a series of images loaded into a QuPath project to a maximum pixel dimension size. We suggest a user-input maximum dimension of 2000 – 3000 pixels, depending on each user’s resolution needs and RAM limitations. The program will save each downsampled image to a folder, including a CSV file containing the downsample scale factor which can be automatically loaded into the PIVOT program during coordinate registration.

### The PIVOT user interface

PIVOT is an open-source program coded in python using the pyQT user-interface development package. This interface enables users to create registration projects, consisting of a single fixed image and one or a series of moving images, to perform various registration tasks. The functions of the user interface are split into five tabs, which we describe separately here. We also generated a user-guide (see supplementary materials) explaining the usage of each tab in extended detail.

Tab 1. Input project settings: this tab allows users to define project settings or to import a previously defined project. A project consists of a single user-defined fixed image (the reference coordinate system for all registration tasks), one or a series of moving images (which will be registered into the fixed image’s coordinate space), and a folder where all job metadata (consisting of the job template file, registration transforms, registered images, and registered coordinate data) will be saved. Once defined, a user may load a previously defined project template file to continue a previous project, incorporate new moving images, or apply registration transforms to coordinate files or high-resolution image files.

Tab 2. Calculate registration of image pairs: this tab allows users to calculated point-cloud and fine-tuned elastic registration of multi-omic image pairs. First, the user will select a moving image to register from a drop-down list of all moving images. The image will load and display side-by-side with the fixed image. Using a series of keyboard shortcuts, the user may rotate, scale, flip, and zoom on the fixed and moving image until similar regions of interest are displayed. If necessary, the user may adjust the brightness and contrast settings of either image to enhance visibility. Next, the user should select a minimum of six fiducial point pairs over similar regions in the fixed and moving image. Once at least six point pairs are selected, point-cloud-based registration may be calculated, optimizing the overlay of the points by transforming their scale, rotation, and translation. After calculation, the interface will display the pre- and post-registered images with calculated RMSE values. If the overlay quality is unacceptable, the user may return to the fiducial selection window. If the overlay quality is acceptable, the user can save these transforms and registered images.

After saving the point-cloud registration transforms, the user has the option to fine-tune the registration through calculation of nonlinear registration using the CODA algorithm.^23^ This fine-tune registration uses maximization of the cross-correlation of image intensity to attempt to correct for local warping between images. For image pairs with similar intensity profiles (such as registration of two brightfield images [H&E to H&E or H&E to IHC] or two fluorescent images, this is straightforward. For image pairs with dissimilar pixel intensity profiles (such as for registration of a brightfield image to a fluorescent image), the program will automatically complement the moving image to improve the performance of the automated registration. For the nonlinear registration calculation, the user may finetune parameters for patch tile size and tile spacing to improve performance. If the elastic registration appears to improve the image overlay, the user may save this transform information. If it does not appear to improve registration, the user may proceed with point-cloud-based affine registration only.

Tab 3. Align coordinate data using calculated registration transforms: this tab allows users to register coordinate data by applying the registration transforms calculated using the multi-omic image pairs. Examples of coordinate data that users may wish to register include per-cell protein expression information from spatial proteomic assays and per-spot coordinates from spatial transcriptomics assays. The tab allows users to import a coordinate file, assign which moving image the coordinates correspond to, and input the columns of the CSV file containing the X and Y coordinates. For example, for spot outputs from the 10x Genomics Visium platform, the X and Y coordinates are in columns ‘E’ and ‘F’, or ‘5’ and ‘6’ (the table accepts alphabetic or numerical inputs). Once these variables are assigned, the interface allows users to load the fixed image, moving image, and coordinate points. The points will be overlaid on the unregistered moving image, the registered moving image, and the fixed image. The user must confirm that the registration applies to the coordinates as expected, after which the registered points may be exported. The exported CSV file will mirror the format of the input CSV, retaining any columns containing associated information for transcriptomic spots or protein expression, and replacing the unregistered coordinate values with the updated registered values. The user may save the registered points in the point-cloud registered space or in the fine-tuned elastic registered space.

Tab 4. Align high-resolution images using calculated registration transforms: this tab allows users to register high-resolution images by applying the registration transforms calculated using the low-resolution image pairs. Examples of high-resolution images that users may wish to register include segmentation masks for brightfield or immunofluorescent images. This tab allows users to import a list of image files and assign the corresponding moving image from the PIVOT project. The program will automatically calculate he scale between each high-resolution image and the lower-resolution image used for registration, and will load and register each file using nearest neighbor interpolation so as not to affect the precision of pixel-level labels in image masks.

Tab 5. View job status: this tab allows users to view the status of each pair of images in the current job. The tab consists of a large table with a row for each moving image. Columns include the number of fiducial point pairs clicked per image, the RMSE calculated following point-cloud registration, the RMSE calculated following the fine-tuning elastic registration, and confirmation of whether or not the resulting transforms have been used to align coordinate data. There are no user tasks on this tab. Instead, this tab allows users to rapidly determine for which image pairs registration has been calculated or needs to be calculated.

### Deconvolution of cellular and expression data into spatial transcriptomics spots

While too specific of an application for incorporation into the user-interface, we consider grouping of cell counts, protein expression, and other coordinate resolved biological signals into spatial transcriptomics spots to be a major use enabled by this software. Expanding on the method originally developed for deconvolution of Visium spatial transcriptomics spots using cell types labelled using segmentation of H&E images,^18^ here we show that additional molecular information may be grouped into transcriptomic spot data from registered CSV coordinate data. In the PIVOT GitHub repository linked in this manuscript, we provide sample code which may be adapted to users’ specific applications.

### Comparison of the effect of time spent annotating to registration accuracy

To determine the effect of poor fiducial placement on registration accuracy, we performed time trials on a subset of the images. For each of the three datasets, we selected a single pair of images for method comparison. From dataset A we chose the high-resolution H&E image and one of the IMC ROIs. From dataset B we chose the high-resolution H&E image and one of the cyclic IF images. From dataset C we chose the Visium-associated H&E image and the mass spectrometry image. For each image pair from the three datasets we meticulously generated 20 pairs of high-accuracy fiducial points per fixed and moving image to serve as validation of method performances. For each image pair, two users additionally performed registration nine times: three times repeated allowing 2 minutes for fiducial pair placement, three times repeated allowing one minute for fiducial pair placement, and three times repeated allowing thirty seconds for fiducial pair placement. We applied the automated nonlinear registration to compare the results of affine and nonlinear registration. Following image registration, we applied the registration transforms to the twenty pairs of validation coordinates to compare accuracy. We plotted the root-mean-squared-error (RMSE) of the validation coordinates in bar graphs.

### Comparison to existing registration workflows

We compared the performance of PIVOT to two existing methods, CODA automated registration, and ImageJ fiducial-point registration to these datasets by comparing root-mean-squared-error (RMSE) and runtime. For each image pair from the three datasets we meticulously generated 20 pairs of high-accuracy fiducial points per fixed and moving image to serve as validation of method performances. We applied PIVOT semi-automated registration, CODA automated registration, and ImageJ fiducial-point registration to all images in the datasets, and calculated root-mean-squared-error (RMSE) and runtime. These data were plotted in a bar plot for comparison.

Application of ImageJ fiducial-point registration was performed using the following steps. Reference and moving image pairs were loaded into ImageJ. For each pair, three corresponding landmarks were manually placed using the multi-point tool. An affine transformation matrix was then computed using the Landmark Correspondences plugin with the following parameters: alpha = 1.00, mesh resolution = 32, and transformation class set to affine. The output affine matrix was exported and loaded into Python. Images were transformed with OpenCV’s warpAffine function. Target points were transformed using numpy. RMSE values were computed for the transformed landmarks to quantify alignment accuracy for each image pair.

